# Cross-link assisted spatial proteomics to map sub-organelle proteomes and membrane protein topology

**DOI:** 10.1101/2022.05.05.490733

**Authors:** Ying Zhu, Kerem Can Akkaya, Diogo Borges Lima, Cong Wang, Martin Lehmann, Fan Liu

## Abstract

The specific functions of cellular organelles and sub-compartments depend on their protein content, which can be characterized by spatial proteomics approaches. However, many spatial proteomics methods are limited in their ability to resolve organellar sub-compartments, profile multiple sub-compartments in parallel, and/or characterize membrane-associated proteomes. Here, we develop a cross-link assisted spatial proteomics (CLASP) strategy that addresses these shortcomings. Using human mitochondria as a model system, we show that CLASP can elucidate spatial proteomes of all mitochondrial sub-compartments and provide topological insight into the mitochondrial membrane proteome in a single experiment. Biochemical and imaging-based follow-up studies demonstrate that CLASP allows discovering mitochondria-associated proteins and revising previous protein sub-compartment localization and membrane topology data. This study extends the scope of cross-linking mass spectrometry beyond protein structure and interaction analysis towards spatial proteomics, establishes a method for concomitant profiling of sub-organelle and membrane proteomes, and provides a resource for mitochondrial spatial biology.

## Introduction

Cellular processes are mediated through complex interactions of biological molecules. To efficiently execute and precisely control interactions and their biochemical reactions in the three-dimensional space, cells are compartmentalized into various membrane-bound and non-membrane-bound compartments that carry out specialized functions. Understanding the spatial distribution of proteins and their dynamics provides crucial insights into the molecular basis of compartment-specific cellular functions. To enable protein localization profiling with high throughput and in a system-wide manner, various liquid chromatography mass spectrometry (LC-MS)-based spatial proteomics methods have been developed (Christopher *et al*, 2021; Lundberg & Borner, 2019), including Dynamic Organellar Maps (Borner, 2020), Localisation of Organelle Proteins by Isotope Tagging (LOPIT) (Christoforou *et al*, 2016), and Protein Correlation Profiling (Krahmer *et al*, 2018; Orre *et al*, 2019). However, these methods depend on subcellular fractionation, which limits their spatial resolution because separation of different organelles is often incomplete and organelle sub-compartments cannot be resolved.

Information on sub-compartment-specific protein localization can be obtained by proximity-dependent enzymatic labeling approaches such as APEX, BioID and similar strategies (TurboID, APEX2, etc.) (Choi & Rhee, 2021; Go *et al*, 2021). These methods (referred to as APEX/BioID for the remainder of the paper) rely on fusing a biotinylating enzyme to a protein of known localization or a peptide sequence targeted to a specific sub-compartment, enabling the enzyme-assisted biotin labeling of proximal proteins. Therefore, APEX/BioID methods require multiple experiments to capture different sub-compartments and applying them to characterize membrane-associated proteomes remains challenging. Furthermore, their labeling radius is difficult to control, compromising the spatial resolution. In addition, the required engineering and ectopic expression of the target protein/peptide may introduce artifacts.

A potential yet unexplored alternative to fractionation- and proximity labeling-based spatial proteomics methods is cross-linking mass spectrometry (XL-MS) (O’Reilly & Rappsilber, 2018). In XL-MS, proteins are covalently linked using small organic molecules (cross-linkers) composed of a spacer arm and two functional groups that are reactive toward specific residues. Subsequently, LC-MS is used to identify residue-to-residue cross-links. A cross-link can only occur if the distance between two residues is small enough to be bridged by the cross-linker. Consequently, the radius of XL-MS is clearly defined by the spacer arm length of the selected cross-linker (Merkley *et al*, 2014), which is typically 5-20 Å (0.5–2 nm). This suggests that XL-MS may enable spatial proteome profiling at a higher and more easily controllable spatial resolution than BioID (ca. 10 nm (Kim *et al*, 2014)), APEX (ca. 20 nm (Martell *et al*, 2012)), and μMap-based labeling (ca. 4 nm and currently limited to cell surface proteins (Geri *et al*, 2020)). However, even though we and others have developed methods for proteome-wide XL-MS (Liu & Heck, 2015; Mendes *et al*, 2019; O’Reilly & Rappsilber, 2018) and have shown that these approaches can capture large parts of the proteome in intact cells and organelles (Fasci *et al*, 2018; Gonzalez-Lozano *et al*, 2020; Jiang *et al*, 2021; Liu *et al*, 2018; Schweppe *et al*, 2017; Wheat *et al*, 2021; Wittig *et al*, 2021), cross-linking has so far only been used to analyze protein structures and interactions.

Here, we demonstrate that, beyond its utility in structural biology and interactomics, cross-linking enables high-resolution systematic mapping of protein localizations. We establish and validate the concept of cross-link assisted spatial proteomics (CLASP) by analyzing intact human mitochondria. We chose mitochondria as a model system because they consist of spatially distinct sub-compartments - the outer membrane (OMM), the intermembrane space (IMS), the inner membrane (IMM), and the matrix (Frey & Mannella, 2000) – allowing us to evaluate whether CLASP is able to (1) determine protein localization with sub-compartment resolution, (2) characterize several sub-compartments in parallel, and (3) capture membrane protein localization. CLASP enables us to determine the specific localizations of 417 proteins across all mitochondria sub-compartments. The CLASP dataset provides sub-mitochondrial localizations for 265 known mitochondrial proteins and 143 proteins previously not assigned to this organelle, and identifies 9 mitochondria-associated cytosolic proteins. Furthermore, the spatial resolution of CLASP is high enough to give insights into the topology of 89 membrane proteins, including 53 proteins for which topological information so far has been incorrect or lacking. We confirm several of these findings through biochemical and imaging approaches, demonstrating the effectiveness and robustness of CLASP for elucidating protein localizations with high spatial resolution.

## Results

### CLASP enables detailed protein localization mapping based on routine XL-MS

CLASP is based on the idea that system-wide XL-MS experiments are very likely to capture some well characterized proteins with known subcellular localization, since such “localization markers” (LMs) tend to be highly abundant. The cross-links of these LMs allow deducing the relative localization of the directly connected proteins. CLASP analysis can thus extract spatial information from any XL-MS dataset of an enclosed biological system, provided that (i) cross-links are formed with a defined labeling radius, (ii) a high fraction of inter-protein cross-links is detected and (ii) some LMs are captured. To assess whether these three conditions can be fulfilled in a standard proteome-wide XL-MS experiment, we analyzed intact mitochondria isolated from HEK293 cells using DSSO cross-linker (Figure 1A). We identified 8478 cross-links from 1460 proteins at 2% spectrum-level FDR, including 3591 intra-protein and 4887 inter-protein links corresponding to 4090 protein-protein connections (Table S1). Comparing these cross-links to known structural features of human mitochondria (Frey & Mannella, 2000; Perkins *et al*, 1997) shows that DSSO labels protein within a clearly defined radius of 4 nm (Supplementary Note 1). These observations confirm that our XL-MS dataset meets the first and second requirement for CLASP. For all following analyses, we further filtered this dataset by removing proteins lacking inter-protein links or forming small disconnected clusters resulting in a filtered interactome of 3466 connections among 850 proteins.

**Figure 1.**
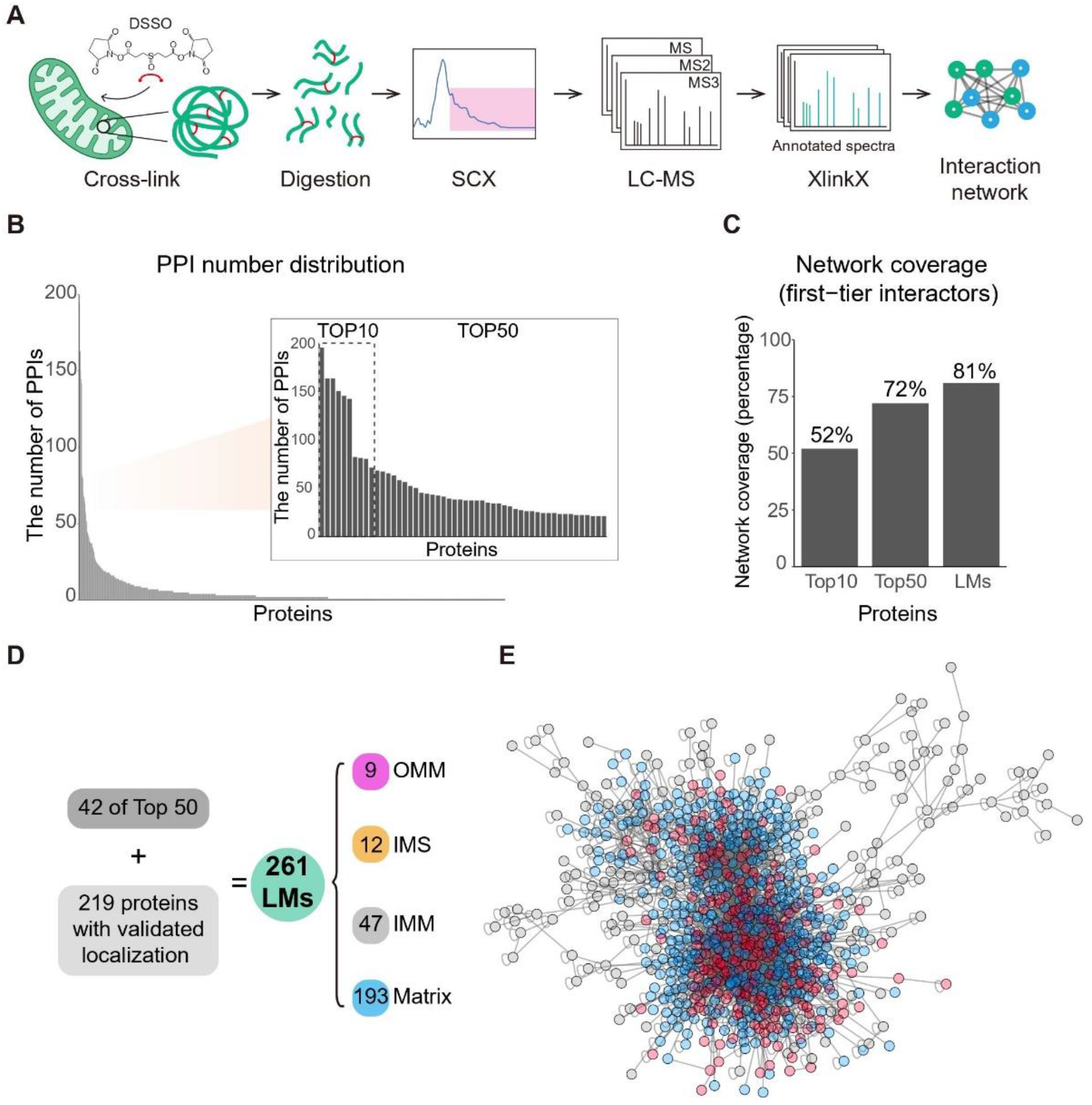
CLASP yields localization information for most proteins in a XL-based mitochondrial PPI network. (A) Workflow for the XL-MS analysis of human mitochondrial proteins. (B) All identified proteins ranked by the number of their interactions. The inset shows the top 10 and top 50 most-connected proteins. (C) Network coverage achieved when considering the first-tier interactors of the top 10 most-connected, top 50 most-connected and all LMs. (D) The origins and subcompartment localizations for all 261 LMs. Proteins were selected as LMs if (i) their sub-mitochondrial localization had been thoroughly established in previous work and (ii) they were part of the top 50 most-connected proteins, had a corresponding PDB structure, or were a component of a well-studied mitochondrial protein assembly. (E) Distribution of LMs (red) and their first-tier interactors (blue) in the mitochondrial PPI network.

To evaluate whether our standard XL-MS experiment also fulfilled the third prerequisite for CLASP – the detection of robust LMs – we assessed the protein connectivity in the XL-based network (Figure 1B). We found that 9 of the 10 most connected proteins and 42 of the 50 most connected proteins have well-established sub-mitochondrial localizations and thus can serve as high-confidence LMs (Table S2). The first-tier interactors of these LMs (i.e. proteins connected through direct cross-links) cover 52% and 72% of the XL-based network, respectively (Figure 1C), confirming that XL-MS readily captures well-connected LMs that make the majority of the detected interactome amenable to CLASP localization predictions.

### CLASP facilitates efficient protein localization mapping

Since our goal is to use CLASP for making biological discoveries with the highest possible confidence, we sought to maximize the number of reliable LMs in our mitochondrial protein network by considering well-characterized mitochondrial proteins and complexes such as VDAC, TIM/TOM complex, mitochondrial contact site and cristae organizing system (MICOS), oxidative phosphorylation complexes (electron transport chain complexes I-IV and ATP synthase) and the mitochondrial ribosome. To assess their suitability as LMs, we curated spatial information from the published literature and high-resolution structures (Table S2). Collectively, we identified 261 high-confidence LMs, including 9 OMM proteins, 12 IMS proteins, 47 IMM proteins, and 193 mitochondrial matrix proteins (Figure 1D, Table S2). Using these LMs, CLASP captured 426 proteins based on first-tier interactions across all mitochondrial sub-compartments (Table S2). These results demonstrate that CLASP allows profiling of multiple sub-compartments in parallel and determining protein localizations with sub-organelle resolution.

We found that 36% (153 proteins) of the CLASP results confirm previous reports, 2% (9 proteins) are ambiguously annotated, 6% (25 proteins) contradict published localization data, and 56% (239 proteins) present new spatial and topological information for previously unannotated proteins(Figure 2A). CLASP suggests new sub-compartment localizations for 164 proteins, revises the topology of 91 membrane proteins (see final Results section for more details), and identifies 9 mitochondrial candidate proteins that have previously not been associated with this organelle.

**Figure 2.**
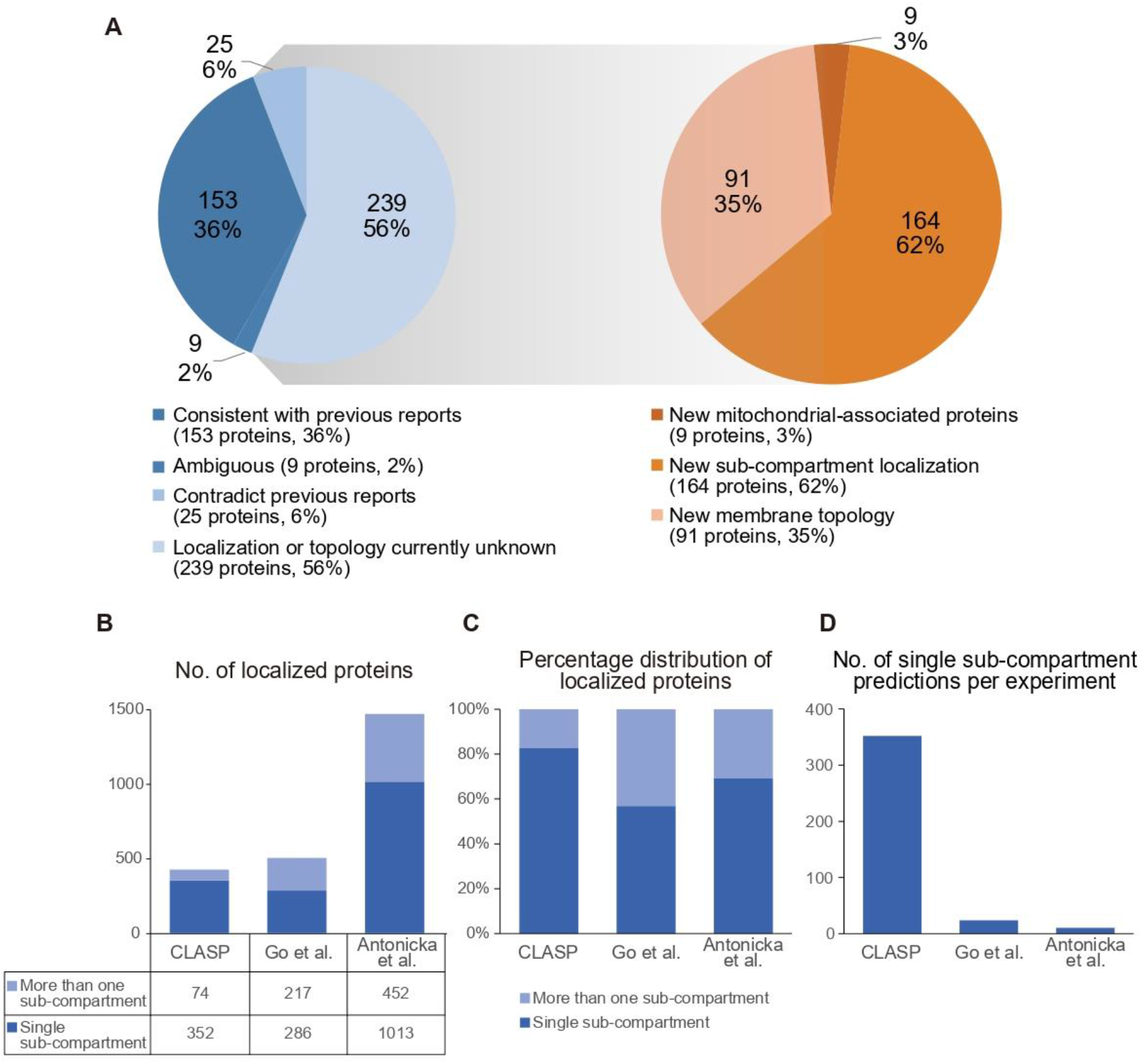
Evaluation of CLASP performance. (A) Comparison of CLASP predictions to published protein localization information (blue pie chart, left) and breakdown of CLASP predictions that disagree with previous reports or relate to previously unannotated proteins (orange pie chart, right). Annotations of individual proteins are shown in Table S2. (**B-C**) Number (B) and (C) percentage distribution of mitochondrial protein localization predictions by CLASP and in published proximity biotinylation resources (Go *et al*., 2021) (Antonicka *et al*., 2020). The columns are sub-divided to indicate the spatial specificity of the localization predictions. (D) Estimated information yield of a single CLASP experiment and single BioID experiments in (Go *et al*., 2021) and (Antonicka *et al*., 2020), calculated as the total number of single sub-compartment predictions divided by the total number of assays.

To further evaluate the scope of CLASP, we compared our dataset to two recently published proximity biotinylation-based spatial proteomics datasets generated in the same cellular system: the HEK293 cell map (Go *et al*., 2021) and the HEK293 mitochondrial proximity interactome (Antonicka *et al*, 2020) (Figure 2B and 2C).

The HEK293 cell map included experiments with 12 mitochondria-specific BioID fusion proteins and determined mitochondrial localizations of 503 proteins by two data analysis algorithms (see Methods), but annotations are limited to three categories (matrix, IMM/IMS and OMM/peroxisome), indicating that the spatial resolution is too low to fully discern all sub-compartments. By contrast, a single CLASP experiments predicted localizations for slightly fewer proteins (426) but resolved 12 categories of sub-compartment localizations (see Table S2, column ‘Subloc’). This allowed us, for example, to identify specific proteins for each individual sub-compartment and distinguish between proteins protruding into different adjacent sub-compartments, e.g. matrix-facing vs. IMS-facing IMM proteins. A similar spatial granularity was reported by Antonicka et al., who also captured the highest number of mitochondrial proteins (1465) (Fig. 2B). However, the fraction of single sub-compartment predictions was lower in the Antonicka et al. dataset than in our CLASP dataset (Fig. 2C), indicating that CLASP predictions provide a higher spatial specificity. Furthermore, achieving this spatial resolution required BioID assays with 100 baits from all mitochondrial sub-compartments (Antonicka *et al*., 2020) but only one CLASP experiment in native mitochondria. This demonstrates that – by obviating exogenous fusion proteins and instead taking advantage of well-characterized mitochondrial proteins to predict new protein localizations – CLASP substantially increases the yield of spatial information from a single experiment (Figure 2D).

### CLASP reveals biologically relevant sub-compartment localizations

Having evaluated the fundamental features of CLASP, we next assessed its potential to provide new biological insights. We predicted sub-compartment localizations for 164 non-membrane proteins that were previously unassigned, mapping 36 proteins to the IMS, 113 to the matrix and 15 to both locales (Table S2). We selected one of these proteins, FAM136A, for complementary validation. The subcellular locale of FAM136A is currently unknown but CLASP predicts that it localizes to the IMS, based on direct connections to 3 IMS LMs and 3 IMM LMs (Figure 3A). To verify this prediction, we transfected a plasmid encoding C-terminally HA-tagged FAM136A into HEK293T and HeLa cells. By confocal and STED microscopy, we found that FAM136A co-localizes with the mitochondrial marker TOMM20 (Figure 3B). By confocal, we found that FAM136A-HA co-localizes with the mitochondrial marker TOMM20. Furthermore, we transfected FAM136A-HA to HeLa-Cox8A-SNAP cells(Stephan *et al*, 2019) and observed that FAM136A-HA formed small clusters within mitochondria by STED microscopy. (Figure 3B). We next checked if FAM136A is soluble or membrane bound by alkaline carbonate extraction. We detected FAM136A only in the soluble fraction, indicating it is not membrane-bound (Figure 3C). Finally, a protease protection assay confirmed the IMS localization of FAM136A (Figure 3D).

**Figure 3.**
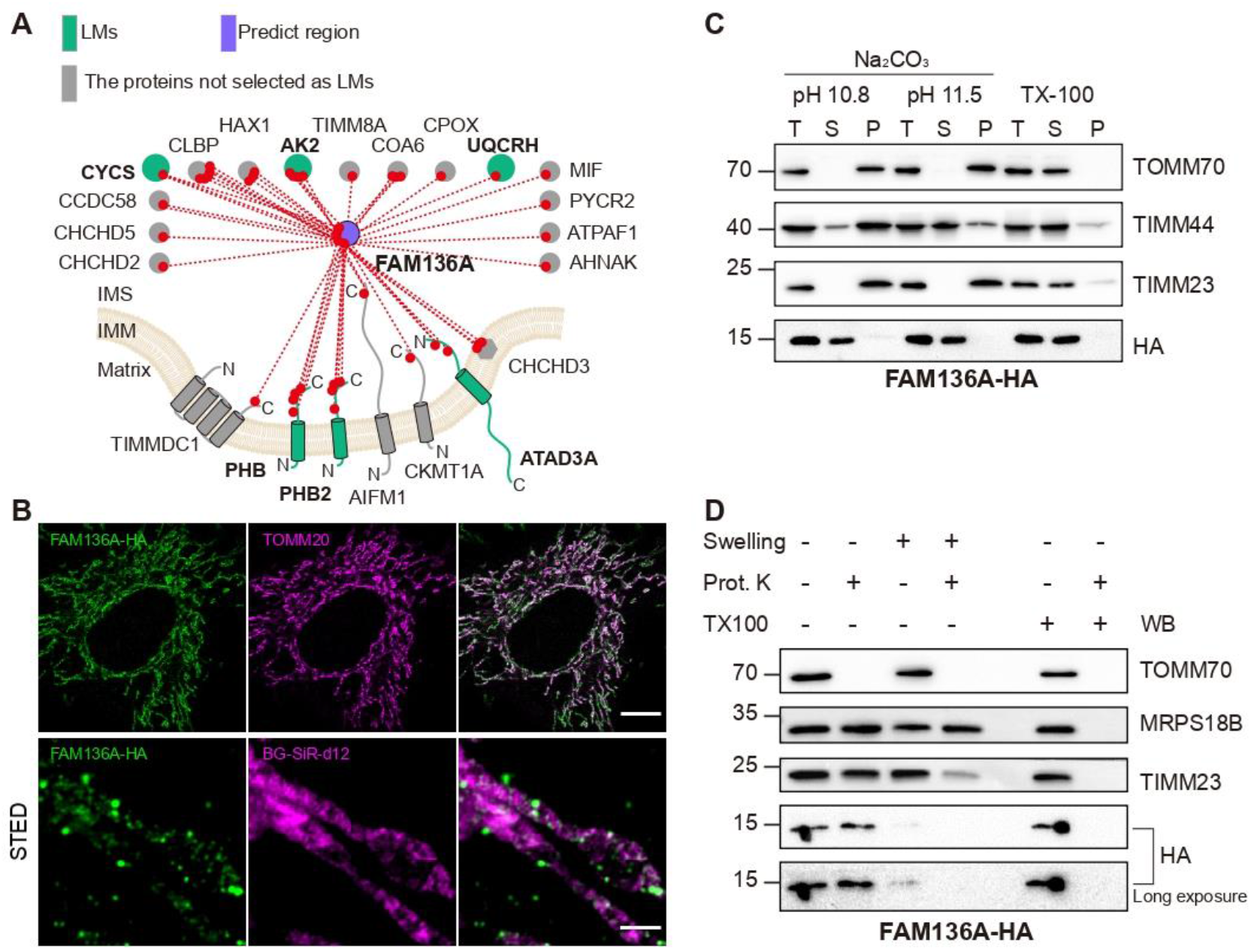
CLASP reveals FAM136A as a new IMS protein. (A) Cross-link map of FAM136A and its interacting proteins. LMs are shown in green; FAM136A is shown in purple; proteins that are not LMs but supporting the CLASP annotation of FAM136A are shown in grey. (B) Confocal fluorescence images of FAM136A-HA, TOMM20 in HeLa cells (upper panel). STED microscopy images in HeLa-COX8A-SNAP cells stained with BG-SiR-d12 and HA antibody (lower panel). Scale bar is 10μm for confocal images and 1μm for STED images. (C) Alkaline carbonate extraction of mitochondria isolated from HEK293T cells overexpressing FAM136A-HA. The OMM protein TOMM70, IMM protein TIMM23 and IMM associated protein TIMM44 are used as markers for each sub-compartment. T, total mitochondrial extraction; S, supernatant; P, pellet of mitochondrial membrane. (D) Protease protection assay combined with WB to analyze the localization of FAM136A-HA. OMM protein TOMM70, IMM protein TIMM23 and matrix protein MRPS18B are used as markers for each mitochondrial sub-compartment.

### CLASP discovers mitochondria-associated proteins

We found 9 proteins, which have no previously reported association with mitochondria, cross-linked to the cytosolic side of OMM LMs (Table S2). Considering the DSSO labeling radius of 4 nm, we hypothesized that these proteins may directly bind to the OMM, potentially at organelle-mitochondria contact sites. One of these proteins, FAF2, is known as an endoplasmic reticulum (ER) protein involved in ER-associated degradation (Mueller *et al*, 2008). We found FAF2 directly connected to the cytosolic parts of two OMM LMs - VADC2 and TOMM5 - as well as the ER- and mitochondria-localized protein CYB5R3 (Neve *et al*, 2012) (Figure 4A), suggesting that FAF2 might localize to both the OMM and the ER.

**Figure 4.**
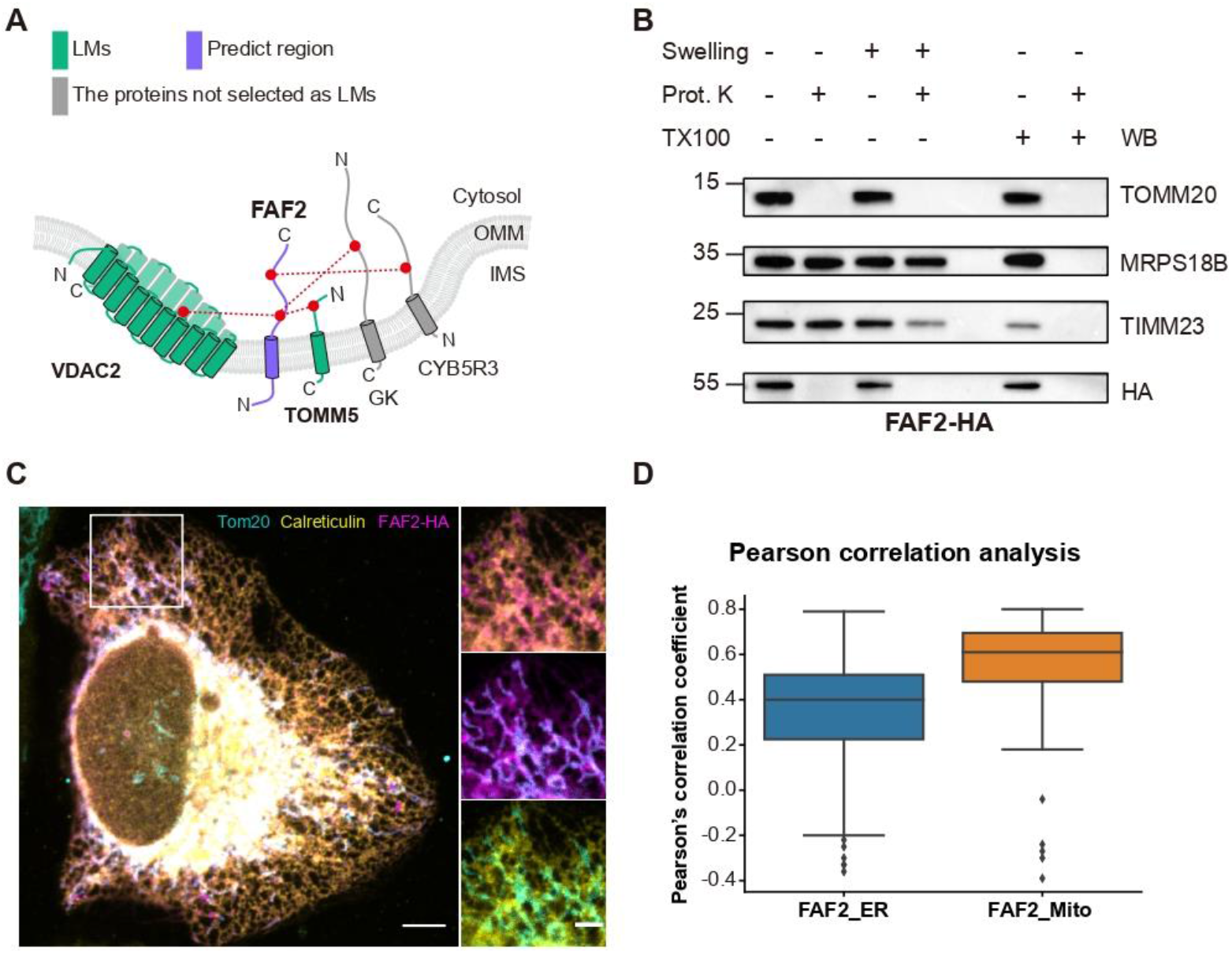
CLASP discovers FAF2 as a new mitochondria-associated protein. (A) Cross-link map of FAF2 and its interacting proteins. LMs are shown in green; FAF2 is shown in purple; proteins that are not LMs but supporting the CLASP annotation of FAF2 are shown in grey. (B) Protease protection assay combined with WB to analyze the localization of FAF2-HA in HeLa cells. OMM protein TOMM70, IMM protein TIMM23 and matrix protein MRPS18B are used as markers for each mitochondrial sub-compartment. (C) Confocal fluorescence images of FAF2-HA, Calreticulin (ER marker) and TOMM20 (mitochondria marker) in HeLa cells. Scale bar is 10μm for the whole cell image and 5μm for the cropped images. Pearson correlation analysis of co-localization of FAF2, ER, and mitochondria. FAF2_ER: FAF2 and ER co-localization signal; FAF2_Mito: FAF2 and mitochondria co-localization signal; n equals to 75 from 4 independent experiments.

To confirm the submitochondrial localization of FAF2, we performed a protease protection assay on purified mitochondria from HEK293T cells (Figure 4B). FAF2 showed a similar digestion pattern as the OMM protein TOMM20, supporting the CLASP prediction that FAF2 is an OMM protein with a cytosol-facing C-terminus. To corroborate the localization of FAF2 at both the ER and mitochondria, we expressed FAF2-HA in HeLa cells and performed confocal imaging of FAF2-HA, OMM marker TOMM20 and ER marker Calreticulin. FAF2 co-localizes with TOMM20 and Calreticulin, indicating a dual localization to both OMM and ER (Figure 4C and 3D). Finally, we employed a SPLICS (split-GFP-based contact site sensors) system (Cieri *et al*, 2018; Vallese *et al*, 2020) to locate ER-mitochondria contact sites. We generated HeLa cells stably expressing OMM-GFP1-10 (OMM targeting sequence fused to GFP fragments containing β-strands 1-10) and ER-GFP11 (ER targeting sequence fused to GFP fragment β-strand 11). We monitored GFP fluorescent signals of contact sites in the presence and absence of FAF2-HA by confocal imaging. FAF2-HA overexpression increased SPLICS positive events compared to the untransfected group, providing further evidence that FAF2 is important for ER-mitochondria contact sites (Figure S1).

### CLASP reveals the topologies of membrane proteins

In the previous sections, we established that CLASP enables high-resolution spatial profiling owing to its clearly defined labeling radius. However, another factor contributing to the high resolution of CLASP is that it provides information on sub-compartment localization on the residue level, i.e. CLASP can potentially distinguish regions within one protein that face different sub-compartments. This offers the opportunity to use CLASP for assessing the topology of membrane proteins. In our dataset, we identified 89 mitochondrial membrane proteins, for which we could assign membrane regions based on previous findings and computational predictions (see Methods for details). For 53 of these proteins, we were able to revise or complete their topological annotation based on CLASP data (Table S2, Figure 5 and Figure S2A). In addition, CLASP suggested topologies for 38 potential membrane proteins with low TM probability in the computational predictions (Figure S2B).

**Figure 5.**
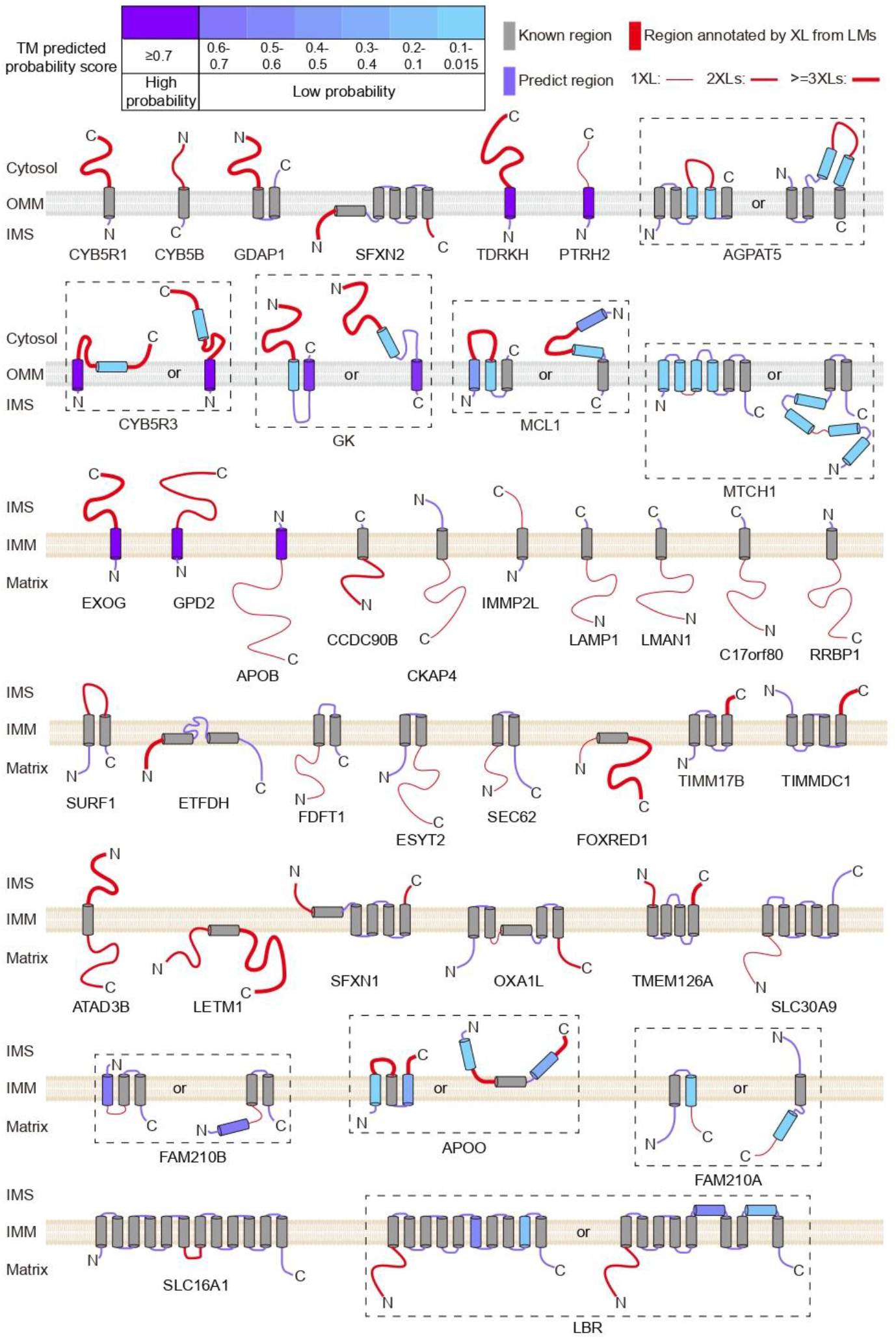
CLASP determines the membrane topologies of 53 OMM and IMM proteins. The TM regions predicted by TMHMM2.0 are gradient colored according to their posterior probabilities of TM helix; Uniprot annotated TM regions are shown in grey; the regions annotated by CLASP are shown in red, the soluble regions predicted by TMHMM2.0 are shown in purple. Of note, TMHMM2.0 cannot predict the localization of the soluble regions, but CLASP allows determining the orientation of the TM regions and localization of the predicted soluble regions. Dashed boxes indicate different possible topologies of one protein. Previously known and predicted TMs are included in Table S2. 41 proteins are presented here and the other 13 proteins are shown in Figure S2A.

To validate these topology predictions, we followed up on CYB5R3 and CYB5B. CLASP predicts that CYB5R3 is an OMM protein with a cytosol-facing C-terminus, while CYB5B is an OMM protein with an IMS-facing C-terminus. We expressed C-terminal HA-tagged versions of each protein in HEK293T cells and performed protease protection assays, confirming their predicted membrane topologies (Figure S3).

Furthermore, we found that CLASP predictions can help correct existing topology annotations. For instance, TMEM126A is an IMM protein with matrix-facing N- and C-termini according to topology information from Swiss-Prot. By contrast, CLASP showed that both termini of TMEM126A cross-link to IMS-localized lysine residues (Figure 6A) and protease protection assays demonstrated that TMEM126A has a similar digestion pattern as the IMM protein COX8A-SNAP, which has an IMS-facing C-terminus (Figure 6B). Both termini are very likely located in the same sub-compartment since TMEM126A has an even number of TM regions. The protease protection data thus confirm the CLASP prediction that the TMEM126 termini protrude into the IMS (Figure 6A, right panel).

**Figure 6.**
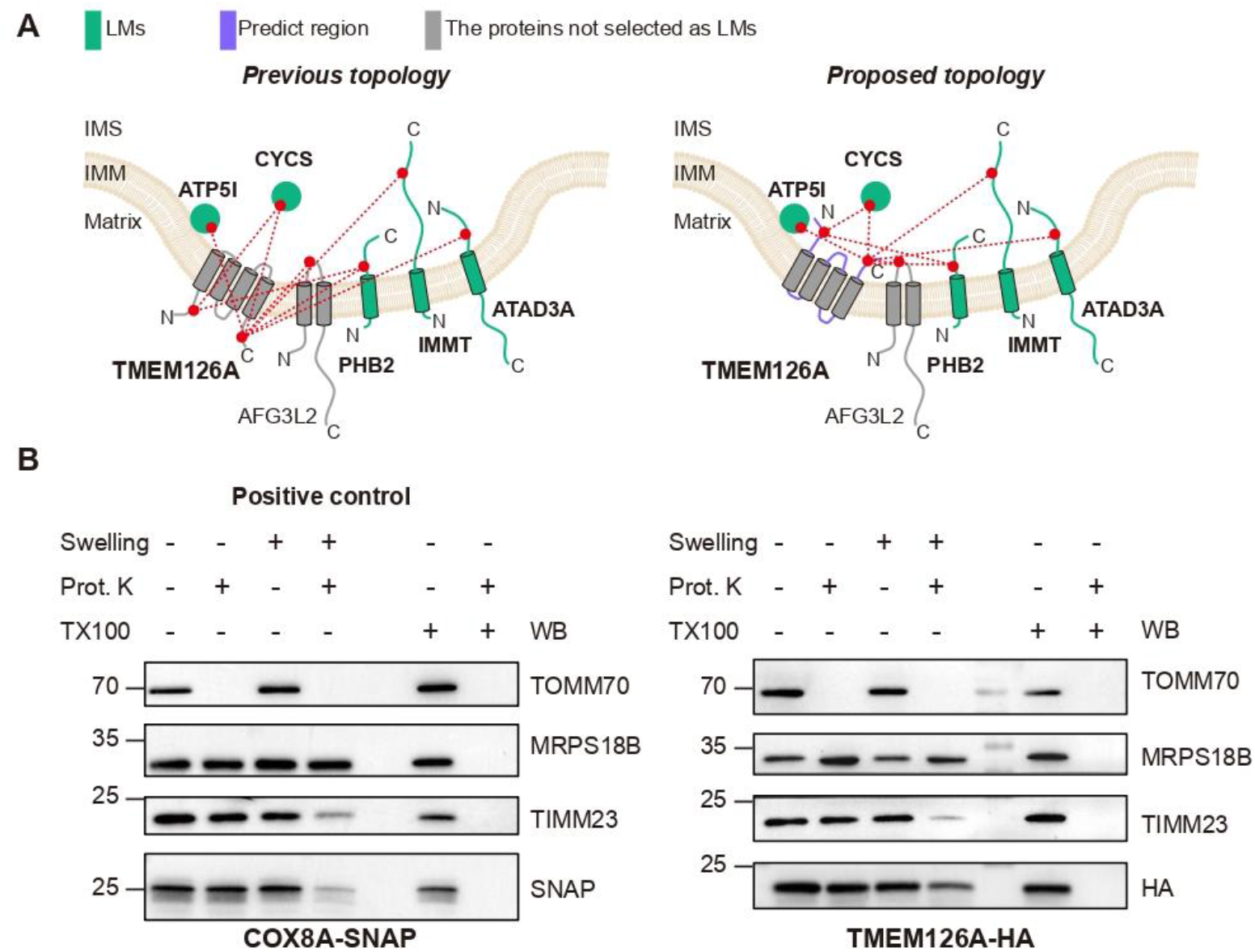
CLASP corrects previously annotated protein TMEM126A. (A) Cross-link map of TMEM126A and its interacting proteins. Left: the cross-link map based on the previous annotation; right: the cross-link map based on the CLASP annotation. LMs are shown in green; predicted regions by TMHMM2.0 are shown in purple; proteins that are not LMs but supporting the CLASP annotation of FAF2 are shown in grey. (B) Protease protection assay combined with WB to analyze the localization of TMEM126A-HA. COXA8-SNAP, which has the same membrane topology as the one CLASP predicted for TMEM126A, serves as a positive control.

To put these findings in context, we compared them to a proximity biotinylation study (Lee *et al*, 2017) that used an APEX approach to predict IMM protein topologies in HEK293 cells (i.e. the same system we used here). Lee et al. designed three APEX fusion-proteins (and several more for validation experiments) to obtain membrane topology information for 60 IMM proteins in the IMM (Figure 7A), whereas CLASP provided topological insights into 67 IMM proteins from a single XL-based mitochondrial protein interaction network. In addition, CLASP allowed us to characterize the topologies of 22 OMM proteins, which could not be analyzed in detail in the experimental setup of Lee et al. This further demonstrates that CLASP is a versatile and efficient strategy to characterize multiple aspects of the spatial proteome in one go.

**Figure 7.**
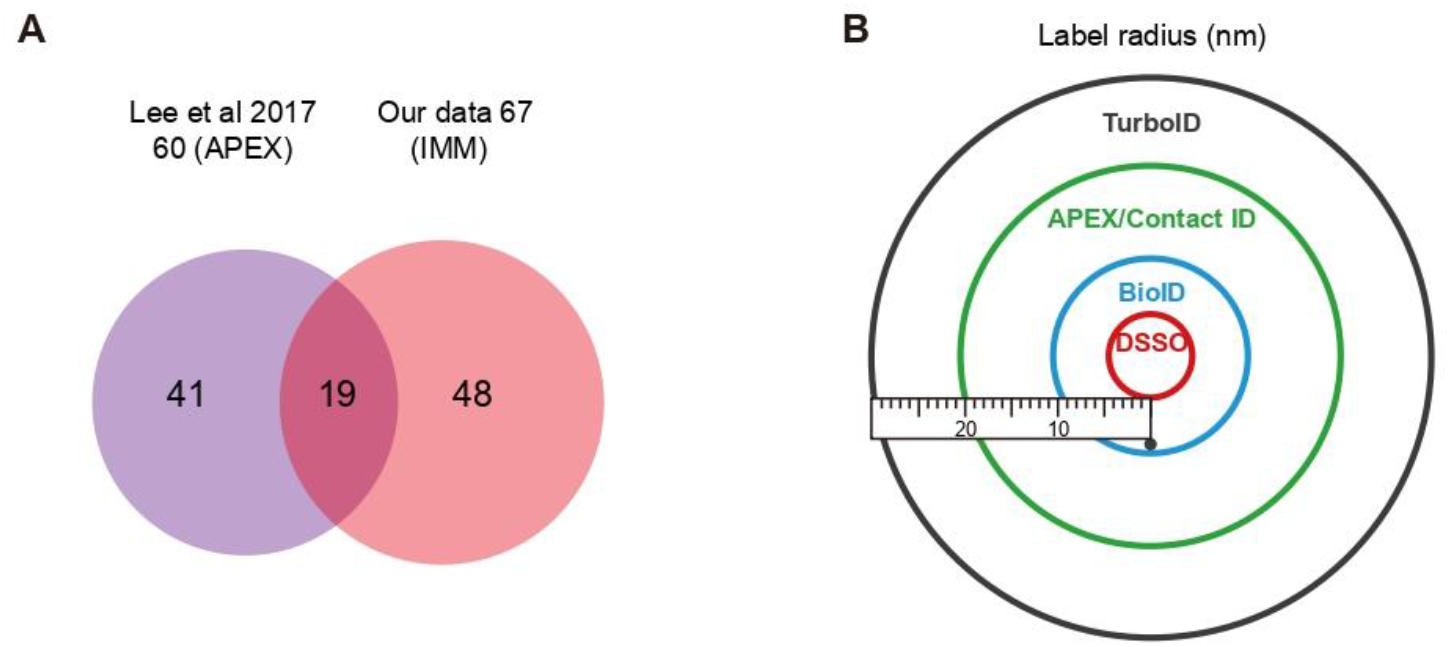
Comparison of CLASP to other proximity labeling-based spatial proteomics methods. (A) Overlap of mitochondrial IMM protein topologies determined by CLASP and in a previous published APEX study (Lee *et al*., 2017). (B) Labeling radii of DSSO-based CLASP and published proximity-labeling methods. DSSO: 4 nm (shown here), BioID: 10nm (Kim et al., 2014), APEX: 20nm (Martell et al., 2012), Contact ID: 10-20nm (Kwak et al., 2020), TurboID: 35nm (May *et al*, 2020)

## Discussion

Over the last two decades, XL-MS has become an established tool in structural biology, complementing methods such as X-ray crystallography and cryoEM with low-resolution structural information on purified proteins and protein complexes in solution. Recent methodological advancements further expanded the scope of XL-MS to more complex biological systems (Liu & Heck, 2015; O’Reilly & Rappsilber, 2018), but XL-MS applications remained focused on the structural analysis of proteins and discovery of protein interactions. In this study, we have developed a framework that extends the application of XL-MS to spatial proteome profiling. Applying purified human mitochondria as a model system, we have shown that CLASP enables simultaneous characterization of the proteomes of multiple sub-compartments, discovery of mitochondria-associated protein, and elucidation of membrane protein topologies in a single experiment. In addition, our analysis suggests that CLASP can provide a more complete picture of the spatial distribution of previously annotated proteins, e.g. for FAF2 (see Figure 4). As another example, our CLASP dataset includes the well-characterized chaperone HSPD1. This protein contributes the correct folding of imported proteins and has previously been annotated as a mitochondrial matrix protein (Cheng *et al*, 1989). Accordingly, CLASP captured connections of HSPD1 to matrix LMs. However, CLASP revealed that HSPD1 is also connected to cytosol-facing regions of the OMM LM TOMM70 and to the two IMS LMs adenylate kinase 2 and cytochrome c (Figure S4). Thus, CLASP indicates that HSPD1 has multiple localization, suggesting that it may be involved in promoting correct folding of imported mitochondrial proteins during the entire translocation process. Previous studies showed that HSPD1 and HSPE1 can be released with cytochrome c and adenylate kinase during apoptosis (Samali *et al*, 1999), which provides evidence for its localization to the IMS and supports our CLASP data. In summary, CLASP allowed us to confirm existing spatial annotation for 153 proteins and add localization and topology data for 264 proteins (Figure 2A). As such, this study not only lays the methodological foundation for future cross-linking-based spatial proteomics studies but also provides a resource for mitochondrial spatial biology.

Considering the above-described insights that can be gained from CLASP analysis, the scope of this approach is most comparable to proximity labeling-based spatial proteomics methods, in particular APEX/BioID-derived strategies. However, a look at previous studies reveals important differences. Several proximity labeling-based spatial proteomics studies also took advantage of the unique morphology of mitochondria and used it as a model system for method benchmarking. These studies demonstrated the capabilities of APEX-based proximity labeling to reveal protein sub-compartment localization (Hung *et al*, 2014; Rhee *et al*, 2013), membrane protein topology (Lee *et al*., 2017), and membrane contact sites (Kwak *et al*, 2020). Characterizing each of these spatial features required the design of dedicated proximity labeling experiments, whereas we have shown that CLASP can provide insights into all these aspects simultaneously in one experiment. Accordingly, CLASP substantially increases the yield of spatial information per experiment (Fig. 2D). The main reason for this fundamental difference is that methods based on BioID or APEX require genetic engineering of fusion proteins targeted to a specific localization. While this strategy has undisputed advantages for profiling cell type- or localization-specific proteomes in organisms (Dumrongprechachan *et al*, 2021; Takano *et al*, 2020; Uezu *et al*, 2016), it requires substantial efforts to generate different constructs and to validate the correct localizations of the fusion proteins. Furthermore, the fusion proteins need to be ectopically expressed, which may affect the cellular status and thereby the spatial distribution of the endogenous proteins-of-interest.

Another important difference between CLASP and proximity-labeling based approaches is the spatial resolution (Figure 7B). The BioID resolution is limited to ca. 10 nm (Kim *et al*., 2014) and, while there have been attempts to reduce the APEX labeling radius (Ke *et al*, 2021), the free diffusion of APEX-generated radicals remains difficult to control, potentially compromising its utility for the highly selective labeling of specific cellular microenvironments. In support of this notion, we provide evidence that, compared to existing proximity labeling resources, CLASP yields a larger fraction of specific localization predictions (Fig 2C). While ambiguous protein localization predictions can be an indicator of biologically relevant dual localizations, e.g. as we observed for HSPD1 (see above and Figure S4), only a method that can confidently predict specific localizations also has the power to reveal potentially relevant dual localizations. For instance, the predictions of (Go *et al*., 2021) do not distinguish between IMM and IMS and can thus not reveal which proteins localize to both or only one of these sub-compartments, whereas CLASP identifies IMS-specific, IMM-specific as well as IMS/IMM-localized proteins.

With the development of CLASP, we introduced the concept of “localization markers” (LMs). CLASP LMs fulfill a similar role as the APEX/BioID fusion proteins, in that both act as “beacons” with well-established localizations that allow deducing the localization of proximal proteins-of-interest. However, the need for engineering and ectopic expression in APEX/BioID-based approaches means that one experiment is generally limited to a single pre-selected “beacon” and thus a single sub-compartment. By contrast, the “beacons”/LMs in CLASP are endogenously present in the sample, and can be selected based on prior knowledge or validation experiments. Being able to consider multiple LMs also increases the localization confidence, since LMs with overlapping interactors allow cross-validating spatial annotations. Many of the localization predictions discussed in this paper, e.g. those for TMEM126A, FAF2, HSPD1 and FAM136A, are indeed supported by the spatial information from at least two LM proteins.

The reliance of CLASP on LMs also means that its predictive power will be lower for biological systems with fewer well-characterized LMs. However, the previous decades of spatial biology research have generated a wealth of protein localization information, yielding hundreds of potential LMs particularly for the most relevant biological systems in fundamental and clinical research. Filling the remaining blank spots in the spatial protein maps of these systems, and thereby revealing new localization patterns of potential biomedical relevance, is one of the main future challenges of spatial biology. CLASP is ideally suited to address this challenge because it can take advantage of existing localization data to reveal new spatial information. At the same time, CLASP is still applicable to systems with sparse endogenous LMs. CLASP predictions in such systems could be improved by ectopically expressing tagged constructs as “exogenous” LMs, which would be conceptually more similar to the APEX/BioID fusion protein approach but still offer the benefits of CLASP’s higher spatial resolution.

In principle, CLASP can use XL-MS datasets of any spatially defined biological system as input, but it is important to note that the prediction power of CLASP critically depends on the comprehensiveness and interconnectivity of the XL-based protein network. Such detailed protein networks can be generated for purified organelles by state-of-the-art XL-MS workflows (Fasci *et al*., 2018; Gonzalez-Lozano *et al*., 2020; Liu *et al*., 2018; Wittig *et al*., 2021), as also shown by the detection depth achieved in this study (8478 cross-links from 1460 proteins). Therefore, CLASP can readily provide detailed localization predictions for any purifiable organelles but would likely be less powerful when applied to intact cells, since proteome coverage of most in-cell XL-MS workflows is currently still limited. However, with recent technological advancements of XL-MS, identification of tens of thousands of cross-links in intact cells is in reach (Jiang *et al*., 2021; Wheat *et al*., 2021) and CLASP paves the way to use these data for elucidating protein localizations across the cell.

## Material and Methods

### Plasmid construction

Total RNA was isolated from HEK293T cells by TRIzol reagent (Invitrogen) according to the manufacturer’s instruction. cDNA library was obtained by using First Strand cDNA Synthesis Kit (Thermo Fisher). For FAF2-HA and CYB5R3-HA, the open reading frames without stop codon were PCR -amplified by Phusion™ High-Fidelity DNA Polymerase (Thermo Fisher) using the forward and reverse primer pairs. The PCR product was cloned into the HindIII and BamHI sites of pcDNA3.1(+) plasmid by using GeneArt™ Seamless Cloning and Assembly Enzyme Mix (Invitrogen). OMP25-HA was cloned into the HindIII and BamHI sites of the pSNAP-N1 plasmid. For SPLICS experiment, ER_GFP11_P2A_OMM_GFP1_10 was PCR-amplified using primer pairs SPICLS-For and SPLICS-Rev. The PCR product was cloned into the Age1 and BamHI sites of pLVX-TetONE-puro plasmid using T4 DNA ligase (Thermo Scientific). CYB5B-HA, FAM136A-HA, TOMM20-Halo-HA and TMEM126A-HA were purchased from Absea Biotechnology Ltd. pSNAPf-Cox8A Control Plasmid was a gift from New England Biolabs & Ana Egana (Addgene plasmid # 101129). The primers used for cloning were synthesized by BioTeZ Berlin-Buch GmbH and are listed in Table S3. All plasmids used in this study were verified by DNA sequencing.

### Cell culture and transfection

HEK293T cells were cultured in Dulbecco’s Modified Eagle Medium (DMEM) supplemented with 10% fetal bovine serum at 37 °C and 10% CO2. Plasmid transfections were performed using Lipofectamine 2000 (Invitrogen) according to the manufacturer’s instruction. Briefly, plasmid DNA and Lipofectamine 2000 were mixed with Opti-MEM separately at a ratio of 3:1 of Lipo2000 to plasmid DNA, and then Lipo2000 was added into the plasmid DNA immediately. The mixture was incubated for 20 min and added dropwise to the cell. After 48 h transfection, cells were washed and harvested with ice-cold PBS for further experiments.

### Lentivirus production via calcium phosphate transfection of HEK293T cells

HEK293T cells were seeded in 6-well plates or in 10 cm petri dishes and were transfected with plasmid mix of packaging plasmid DNA psPAX2, lentiviral envelope plasmid pMD2.G and genomic plasmid DNA (pLVX-TetONE-puro_ER_GFP11_P2A_OMM_GFP1_10) by using calcium phosphate. After 24 h, transfection efficiency was checked with a basic fluorescent microscope. For 6 days, the supernatant was collected and fresh medium was added every 48 h. The supernatant was centrifuged for 5 min at 2,000×rpm to remove cell debris, and filtered using 0.45 μm pore size filters. All supernatants were stored at 4°C for up to 2 weeks or at −80 °C for long-term storage. The concentrated supernatants were collected by centrifugation at 4,696×g for 20 min and stored at −80°C.

### Stable cell line generation for SPLICS system

HeLa cells were seeded at a confluency of 60-70% in 6-well plates or 10 cm petri dishes containing DMEM supplemented with 10% FBS and 1% Pen/Strep. After 24 h, the medium volume was reduced to 1.5 ml for 6-well plates or to 6 ml for 10 cm petri dishes. 0.5-1 ml of non-concentrated or 20-80 μl of concentrated virus was used to infect the seeded cells. After 48-72 h, the control cells were checked with a fluorescence microscope for transduction efficiency indicated by the number of cells expressing free GFP. For the initial selection of successfully infected cells and non-infected cells, up to 2 μg/ml Puromycin were used. Single clones with uniform expression of SPLICS were selected by limited dilution. Additionally, expression of SPLICS system on the ER and mitochondria was verified by expressing either cytosolic GFP11 or GFP1-10 coupled to mCherry, respectively. The cells were continuously checked for viability and expression level of the SPLICS system by addition of 2 μg/mL Doxycycline for 16-20 h of induction and subsequent immunofluorescent staining.

### Mitochondria isolation

Mitochondria isolation was modified from previous published protocols (Wieckowski *et al*, 2009). Briefly, cells were resuspended in ice-cold buffer M (220 mM mannitol, 70 mM sucrose, 5 mM HEPES-KOH pH 7.4, 1 mM EGTA-KOH pH 7.4, supplemented with 1 mM PMSF and complete protease inhibitor EDTA-free cocktail). Cells were lysed by homogenization (25 strokes, 2 times, 900×rpm) using dounce homogenizer. Cell debris were spun down at 800×g for 5 min at 4°C 2 times. The supernatants were centrifuged at 10,000×g for 10 min at 4°C and the pellet was collected. The pellet containing crudely purified mitochondria was further subjected to discontinuous percoll gradient centrifugation (SW41 Ti rotor, Beckman) to obtained high purity mitochondria. Protein concentration was determined by Bradford assay (Bio-rad). The crudely purified mitochondria were used for protease protection assay and alkaline carbonate extraction experiment. High purity mitochondria were used for cross-linking experiment.

### Protease protection assay

Protease protection assays were performed following previous published protocols (Mick *et al*, 2012). Briefly, freshly isolated mitochondria were suspended in SEM buffer (250mM sucrose, 10mM MOPS, 1mM EDTA, pH7.2), EM buffer (10mM MOPS, 1mM EDTA, pH 7.2) or EM buffer containing 1% Triton X-100 respectively and incubated on ice for 25 min. Proteinase K (PK) was added into the samples and incubated for 10 minutes on ice. The reaction was quenched by addition of 2 mM PMSF, followed by trichloroacetic acid (TCA) precipitation. After treatment, the pellet was resuspended in SDS sample buffer and subjected to western blot analysis.

### Alkaline carbonate extraction

Alkaline carbonate extraction was performed following previous published (Mick *et al*., 2012). Briefly, freshly isolated mitochondria were suspended in 0.1 M Na2CO3 pH 11.5, 0.1 M Na2CO3 pH 10.8 or SEM buffer containing 1% Triton X-100 respectively and incubated on ice for 20 min. The samples were centrifuged at 100,000×g for 1 h at 4°C (S55A2 rotor, Thermo fisher). The pellets were resuspended in SDS sample buffer and subjected to western blot analysis.

### Western blot analysis

Protein samples were subjected to SDS-PAGE (14% gel) and wet-transferred to a 0.2 μm Immobilon-PSQ PVDF membrane (Millipore) at 110 V for 90 min. Blots were blocked with 5% BSA for 1 h at room temperature and incubated with corresponding primary antibodies at 4°C overnight. The following antibodies were used in this study: anti-HA (1:1000, mouse, Abcam, ab18181); anti-SNAP (1:1000, rabbit, NEB, P9310S); anti-MRPS18B, anti-TIMM44 and TOMM70 (1:500, 1:1000, 1:500, rabbit, ProteinTech, 16139-1-AP, 13859-1-AP, 14528-1-AP); anti-TIMM23 (1:1000, mouse, DB Biotech, 611222). After washing 3 times with TBST, blots were incubated with secondary antibody peroxidase-conjugated affinipure goat anti-mouse lgG and peroxidase-conjugated affinipure goat anti-rabbit lgG (H+L, Jackson ImmunoResearch, 115-035-003, 111-035-144) for 1 h at room temperature. After washing 3 times with TBST, blots were developed with Pierce ECL Western Blotting Substrate (Thermo Fisher) and imaged by ChemiDoc MP Imager (Bio-Rad).

### Cross-linking of mitochondria

The mitochondrial pellet was resuspended to 1 mg/ml in Buffer M and cross-linked twice with 0.5 mM disuccinimidyl sulfoxide (DSSO), each for 20 min at room temperature with constant mixing. The reaction was quenched with 20 mM Tris-HCl, pH 8.0 for 30 min at room temperature. Mitochondria were collected by centrifugation at 10000×g for 10 min at 4°C.

### Sample preparation for LC-MS

Cross-linked mitochondria were digested in solution. Briefly, urea was added to the mitochondrial pellet to reach a final concentration of 8 M. Proteins were reduced and alkylated with 5 mM DTT (1 h at 37 °C) and 40 mM chloroacetamide (30 min at room temperature in the dark). Proteins were digested with Lys-C at an enzyme-to-protein ratio of 1:75 (w/w) for 4 h at 37 °C. After diluting with 50 mM TEAB to a final concentration of 2 M urea, trypsin was added at an enzyme-to-protein ratio of 1:100 (w/w) for overnight at 37 °C. The digestion was quenched by adding formic acid to a final concentration of 1%. Peptides were desalted with Sep-Pak C18 cartridges (Waters) according to the manufacturer’s protocol, dried under SpeedVac.

### Strong cation exchange (SCX) fractionation of cross-linked peptides

Strong cation exchange (SCX) fractionation was performed on digested peptides using PolySULFOETHYL-ATM column (100 × 4.6 mm, 3 μm particles, PolyLC INC.) on an Agilent 1260 Infinity II UPLC system. A 95 min gradient was applied and fractions were collected every 30 sec. 50 late SCX fractions were desalted by C18 stageTip and dried under SpeedVac.

### LC-MS analysis

Collected SCX fractions were analyzed by LC-MS using an UltiMate 3000 RSLC nano LC system coupled on-line to an Orbitrap Fusion Lumos mass spectrometer (Thermo Fisher Scientific). Reversed phase separation was performed with an in-house packed C18 analytical column (Poroshell 120 EC-C18, 2.7μm, Agilent Technologies). 3 h LC-MS runs were performed for cross-linking acquisition and 2 h runs were done for proteomic analysis. Cross-link acquisition was performed using a CID-MS2-MS3 acquisition method. MS1 and MS2 scans were acquired in the Orbitrap mass analyzer and MS3 scans were acquired in the ion trap mass analyzer. Notably, MS3 acquisitions were only triggered when peak doublets with a specific mass difference (Δ = 31.9721 Da) were detected in the CID -MS2 spectra, as this is indicative for the presence of DSSO cross-linked peptides. The following MS parameters were applied: MS resolution 120,000; MS2 resolution 60,000; charge state 4-8 enable for MS2; MS2 isolation window, 1.6 m/z; MS3 isolation window, 2.5 m/z; MS2-CID normalized collision energy, 25%; and MS3-CID normalized collision energy, 35%. For mitochondrial proteome analysis, mass analysis was performed using an Orbitrap Fusion mass spectrometer (Thermo Fisher Scientific) with a HCD-MS2 acquisition method. MS1 scans were acquired in the Orbitrap mass analyzer and MS2 scans were acquired in the ion trap mass analyzer. The following MS parameters were applied: MS resolution 120,000; MS2 resolution 60,000; charge state 2-4 enable for MS2; MS2 isolation window, 1.6 m/z; MS2-HCD normalized collision energy, 30%.

### MS data analysis

For cross-linking samples, raw data were converted into MGF files in Proteome Discoverer (version 2.4). Data analysis was performed using XlinkX standalone with the following parameters: minimum peptide length 6; maximal peptide length 35; missed cleavages 3; fix modification: Cys carbamidomethyl; variable modification: Met oxidation; DSSO cross-linker =158.0038 Da (short arm = 54.0106 Da, long arm = 85.9824 Da); precursor mass tolerance 10 ppm; fragment mass tolerance 20 ppm. MS2 spectra were searched against a reduced target-decoy UniProt human database derived from proteins combining MitoCarta2.0 database and protein identification from a mitochondrial proteomic measurement. Results were reported at 2% FDR at CSM level. Protein-protein interaction (PPI) network was constructed by Cytoscape software (version 3.8.2). Self-links and small clusters were removed and the main cluster containing 850 proteins (3466 PPIs) was used for CLASP analysis.

Proteomics data were analyzed using MaxQuant software (version 1.6.2.6a) with the following searching parameters: precursor mass tolerance 20 ppm, fragment mass tolerance 20 ppm; fixed modification: Cys carbamidomethylation; variable modification: Met oxidation, protein N-term Acetyl; enzymatic digestion: trypsin/P; maximum missed cleavages: 2. Database search was performed using Swiss-Prot database of all human proteins without isoform (retrieved on May 2020, containing 20,365 target sequences); false discovery rate: 1%. Intensity-based absolute quantification (iBAQ) was enabled to determine to relative abundances of proteins.

### Structural validation for cross-links

Cross-links were mapped onto selected high-resolution structures using Pymol 2.1.0 (Schrodinger LLC). If homologous non-human structures were used, sequences were aligned to the human protein sequence using NCBI BLAST and cross-links were mapped to the aligned residues. The following PDBs were used: the mitochondrial electron transport chain complexes I, II, III and IV (PDB: 5XTD, 1ZOY, 5XTE and 5Z62), TOMM complex (7CK6), TIMM22 complex (7CGP), TIMM9-TIMM10 complex (2BSK), succinyl-CoA ligase complex SUCLG1-SUCLG2 (6G4Q), MCAD-ETF complex (1T9G), 39S mitoribosome (7OIE), frataxin bound iron sulfur cluster assembly complex (6NZU), iron sulfur cluster assembly (5KZ5), calcium uniporter homocomplex (6WDN), transcription ignition complex (6ERQ), mitochondrial DNA replicase (4ZTU), trifunction protein (6DV2), FIS (1PC2), CYB5R3 (1UMK), TIMM44 (2CW9), MUT (3BIC), VDAC1 (6G6U), VDAC2 (4BUM), SLC25A13 (4P5W), COQ8A (4PED), EXOG (5T5C), CLPP (6DL7), PHB2 (6IQE), OPA1 (6JTG), TRAP1 (4Z1L), CKMT1A (1QK1), PNPase (5FZ6).

### Confocal immunofluorescence and STED microscopy

HeLa cells and HeLa-COX8A-SNAP cells (Stephan *et al*., 2019) were seeded on μ-Slide 8 Well Glass Bottom (ibidi, REF 80827) one day before transfection, cultivated in DMEM containing 10 % FBS, 1 % P/S and 1% L-Glut. After 24 h, the transfected cells were fixed with PBS containing 4% paraformaldehyde (PFA) and 4% sucrose for 10 minutes at room temperature. Afterwards, fixation was quenched by removing PFA and adding PBS containing 0.1 M glycine and 0.1 M NH4Cl for 10 min at room temperature. Cells were permeabilized by PBS containing 0.15 % TritonX-100 for 10 min at room temperature and washed by PBS two times. Then sequentially incubated in blocking buffer (PBS containing 1%BSA and 6% NGS) for 30 min at room temperature, blocking buffer containing primary antibodies for 1 h at room temperature. After three washes in blocking buffer, the cells were incubated in blocking buffer containing secondary antibody for 30 min at room temperature. After three washes in blocking buffer, the cells were imaged on a Nikon spinning disc microscope (Yokogawa spinning disk CSU-X1) equipped with the following lasers (488-nm, 561-nm and 638-nm), a 60 × oil objective (Plan-Apo, NA 1.40 Nikon), a 40 × air objective (Plan Apo, NA 0.95), an Andor camera (AU888, 13 μm/pixel) and NIS software. STED images were taken on a STEDYCON system (Abberior Instruments) mounted onto a Nikon TI Eclipse with a nanoZ Controller (Prior) and controlledby Micromanager. Imaging was done with two different pulse Diode lasers (561 nm, 640 nm) for excitation paired with two single counting avalanche photodiodes (650 - 700 nm, 575 – 625 nm) for detection, a 775 nm STED laser for depletion and an optic tunable filter to modulate all laser beams for confocal imaging. Images were captured by using an 100x 1.45 NA lambda oil objective lens with a fixed pixel size set to 20 nm for STED images.

The following primary antibodies were used in FAF2 experiment: anti-mouse-TOMM20 (1:100, Santa Cruz, sc-17764), anti-rat-HA (1:100, Chromotek, 7c9-100), anti-rabbit-Calreticulin (1:100, Thermo, PA3-900). The primary antibodies were used in FAM136A experiment: anti-mouse-TOMM20 (1:100, Santa Cruz, sc-17764), anti-rabbit-HA (1:100, Cayman, cay162200-1). The following secondary antibodies were used in FAF2 experiment: anti-rat-AF594 (1:200, Invitrogen, A11007) and anti-mouse-AF594 (1:200, Thermo, A11032). The following secondary antibodies were used in FAM136A experiment: anti-rabbit-AF647 (1:200, Thermo, A21244) and anti-mouse-AF594 (1:200, Thermo, A11032).

### Imaging analysis

Images were processed using FIJI (https://imagej.net/software/fiji/) image analysis software. For the Pearson’s correlation analysis, a dataset consisting of twoconditions over four experiments with a total of 150 images was used. Cells were manually cropped for measuring the cell-wise Pearson’s correlation coefficient above threshold with a Coloc2 (https://imagej.net/plugins/coloc-2) based script. A bisection-based threshold regression with a PSF of sigma=3pixel was applied.

For the analysis of the SPLICS system, a dataset consisting of 2 conditions over 7 experiments with a total of 336 images acquired as z-stacks with a 0.3 μm step size over 25 slices was used. Analysis was performed on manually cropped out single cells, before the z-stack was subjected to max intensity projection and background subtraction. A Gaussian filter with a sigma of 2 pixel was used to further filter the processed images before applying the Spot Counter plugin (https://imagej.net/plugins/spotcounter) with a box size of 10 and a noise level set to 1,000 to identify and count the SPLICS positive spots.

### Computational predictions for TM regions

For the 426 first-tier interactors of the 261 LMs, we used the software TMHMM (Krogh *et al*, 2001) to predict the transmembrane regions. Here we used the posterior probability to show the prediction confidence. 56 transmembrane proteins with known topology in 261 LMs were used to define the probability score range. Posterior probability score above 0.7 was defined as high confidence transmembrane region, 0.015-0.7 as potential transmembrane region, below 0.015 as soluble region.

### Comparison of CLASP data to published proximity biotinylation datasets

We re-analyzed data from the HEK293 cell map (Go *et al*., 2021), the HEK293 mitochondrial proximity interactome (Antonicka *et al*., 2020) and the IMM protein architecture map (Lee *et al*., 2017).

Our analysis of the Go et al. dataset was based on Supplementary Table 9 of the original paper (Go *et al*., 2021), which shows cellular sub-compartment predictions for the confidently identified proximity interactors (filtered at 1% Bayesian FDR) based on two prediction algorithms – Non-negative Matrix Factorization (NMF) and Spatial Analysis of Functional Enrichment (SAFE). We filtered for proteins for which both algorithms suggested a mitochondrial localization. The number of predicted sub-compartments was determined using NMF predictions since SAFE did not provide any single sub-compartment predictions.

For the Antonicka et al. dataset, we determined the number of confidently predicted proteins by filtering their full proximity interactome (Table S4 in (Antonicka *et al*., 2020)) at a Bayesian false discovery rate threshold of 1% (i.e. the same cutoff as in the original publication). We then used the reported sub-compartment annotations of each BioID bait (Figure 2B in (Antonicka *et al*., 2020)) to determine in which sub-compartments the reported prey proteins were detected.

For the comparison shown in Fig 2B-D, proteins predicted to reside in a single sub-compartment were classified in “single sub-compartment” group. Proteins with ambiguous localization or predicted to reside in more than one subcompartments were classified in “more than one sub-compartments” group (see Table S2, column “single sub-compartment (Yes or No)”).

For the Lee et al. dataset, we considered all transmembrane IMM proteins, for which the authors proposed or confirmed topological information (reported in Data Set S7 of (Lee *et al*., 2017)).

## Acknowledgements

The work was supported by Deutsche Forschungsgemeinschaft Grant (DFG) SFB 958(Z03) to YZ, the Leibniz-Wettbewerb (K284/2019) to CW and Leibniz Association to KA and DBL.

## Author contributions

YZ performed the most of the experiments and data analysis. KA performed the confocal and STED imaging experiments. DBL, CW, and ML assisted the experiments and data analysis. YZ and FL wrote the manuscript. FL developed the concept and supervised the research. All authors reviewed and edited the manuscript.

